# Hypoxia-inducible factor 1α is required to establish the larval glycolytic program in *Drosophila melanogaster*

**DOI:** 10.1101/2025.01.07.631819

**Authors:** Yasaman Heidarian, Tess D. Fasteen, Liam Mungcal, Kasun Buddika, Nader H. Mahmoudzadeh, Travis Nemkov, Angelo D’Alessandro, Jason M. Tennessen

**Affiliations:** Department of Biology, Indiana University, Bloomington, IN 47405, USA; Department of Biochemistry and Molecular Genetics, Anschutz Medical Campus, University of Colorado School of Medicine, Aurora, CO 80045, USA; Affiliate Member, Melvin and Bren Simon Cancer Center, Indianapolis, IN, 46202, USA

**Keywords:** *Drosophila melanogaster*, Hypoxia-inducible factor 1α, Estrogen Related Receptor, glycolysis, Warburg Effect

## Abstract

The rapid growth that occurs during *Drosophila* larval development requires a dramatic rewiring of central carbon metabolism to support biosynthesis. Larvae achieve this metabolic state, in part, by coordinately up-regulating the expression of genes involved in carbohydrate metabolism. The resulting metabolic program exhibits hallmark characteristics of aerobic glycolysis and establishes a physiological state that supports growth. To date, the only factor known to activate the larval glycolytic program is the *Drosophila* Estrogen-Related Receptor (dERR). However, dERR is dynamically regulated during the onset of this metabolic switch, indicating that other factors must be involved. Here we discover that Sima, the *Drosophila* ortholog of Hif1α, is also essential for establishing the larval glycolytic program. Using a multi-omics approach, we demonstrate that *sima* mutants fail to properly activate aerobic glycolysis and die during larval development with metabolic defects that phenocopy *dERR* mutants. Moreover, we demonstrate that dERR and Sima/Hif1α protein accumulation is mutually dependent, as loss of either transcription factor results in decreased abundance of the other protein. Considering that the mammalian homologs of ERR and Hif1α also cooperatively regulate aerobic glycolysis in cancer cells, our findings establish the fly as a powerful genetic model for studying the interaction between these two key metabolic regulators.

**STRUCTURED ABSTRACT:** *Objectives:* The rapid growth that occurs during *Drosophila* larval development requires a dramatic rewiring of central carbon metabolism to support biosynthesis. Larvae achieve this metabolic state, in part, by coordinately up-regulating the expression of genes involved in carbohydrate metabolism. The resulting metabolic program exhibits hallmark characteristics of aerobic glycolysis and establishes a physiological state that supports growth. To date, the only factor known to activate the larval glycolytic program is the *Drosophila* Estrogen-Related Receptor (dERR). However, dERR is dynamically regulated during the onset of this metabolic switch, indicating that other factors must be involved. Here we examine the possibility the *Drosophila* ortholog of Hypoxia inducible factor 1α (Hif1α) is also required to activate the larval glycolytic program.

*Methods:* CRISPR/Cas9 was used to generate new loss-of-function alleles in the *Drosophila* gene *similar* (*sima*), which encodes the sole fly ortholog of Hif1α. The resulting mutant strains were analyzed using a combination of metabolomics and RNAseq for defects in carbohydrate metabolism.

*Results:* Our studies reveal that *sima* mutants fail to activate aerobic glycolysis and die during larval development with metabolic phenotypes that mimic those displayed by *dERR* mutants. Moreover, we demonstrate that dERR and Sima/Hif1α protein accumulation is mutually dependent, as loss of either transcription factor results in decreased abundance the other protein.

*Conclusions:* These findings demonstrate that Sima/HIF1α is required during embryogenesis to coordinately up-regulate carbohydrate metabolism in preparation for larval growth. Notably, our study also reveals that the Sima-dependent gene expression profile shares considerable overlap with that observed in *dERR* mutant, suggesting that Sima/HIF1α and dERR cooperatively regulate embryonic and larval glycolytic gene expression.

**HIGHLIGHTS:**

- The *Drosophila melanogaster* gene *similar* (*sima*), which encodes the sole fly ortholog of Hif1α, is required to up-regulate glycolysis in preparation for larval growth.
- *sima* mutant larvae exhibit severe defects in carbohydrate metabolism and die during the second larval instar.
- *sima* mutant larvae exhibit the same metabolic phenotypes as *Drosophila Estrogen Related Receptor* (*dERR*) mutants, suggesting that these two transcription factors coordinately regulate the larval glycolytic program.
- Sima/Hif1α and dERR accumulation is mutually dependent, as loss of either transcription factor results in decreased abundance of the other.

## INTRODUCTION

When reared under ideal laboratory conditions, larvae of the fruit fly *Drosophila melanogaster* increase their body mass by ∼200-fold in four days (Church and Robertson, 1966). This remarkable growth rate requires that dietary nutrients are efficiently utilized for both energy generation and biosynthesis (Gillette et al., 2021, Chatterjee and Perrimon, 2021). Previous studies revealed that this rapid growth phase is made possible, in part, by an embryonic metabolic switch, which results in the transcriptional up-regulation of genes involved in carbohydrate metabolism, including those that encode enzymes involved in glycolysis and the pentose phosphate pathway, as well as Lactate dehydrogenase (Tennessen et al., 2011, Tennessen et al., 2014b). The resulting metabolic program exhibits the hallmark characteristics of aerobic glycolysis and establishes a physiological state that is ideally suited for biomass accumulation.

The predictable manner by which larval carbohydrate metabolism is coordinately activated establishes the fly as a powerful model to dissect the *in vivo* molecular mechanisms that initiate aerobic glycolysis. In this regard, previous studies of larval glycolytic metabolism focused on the *Drosophila* Estrogen-Related Receptor (dERR), which represents the sole fly ortholog of the ERR family of nuclear receptors and functions as an essential regulator of aerobic glycolysis (Ostberg et al., 2003, Tennessen et al., 2011). During the latter half of embryogenesis, dERR directly activates the expression of genes involved in carbohydrate metabolism, thereby establishing the larval metabolic state. In *dERR* mutants, this metabolic switch fails, and the transcription of genes involved in glycolysis and other aspects of carbohydrate metabolism remain at a basal expression level (Tennessen et al., 2011). As a result, *dERR* mutant larvae are unable to utilize dietary carbohydrates and ultimately die during the mid-L2 larval stage.

While dERR has proven to be a valuable focal point for understanding how glycolysis is coordinately regulated with growth and development, other factors must be involved in the embryonic metabolic switch. In this regard, the *Drosophila* gene *similar* (*sima*), which encodes the sole fly homolog of the Hypoxia Inducible Factor 1α (Hif1α) (Nambu et al., 1996, Lavista-Llanos et al., 2002, Bacon et al., 1998), represents an intriguing candidate gene for regulating glycolytic gene expression. Throughout the animal kingdom, Hif1α regulates gene expression in response to changes in oxygen availability (Kaelin and Ratcliffe, 2008, Loenarz et al., 2011). Under normoxic conditions, Hif1α is hydroxylated in an oxygen- and α-ketoglutarate-dependent manner by the enzyme prolyl hydroxylase 2 (PHD2), ubiquitinated by a complex that includes VHL, and subsequently degraded (Epstein et al., 2001, Bruick and McKnight, 2001, Cockman et al., 2000, Kamura et al., 2000, Tanimoto et al., 2000, Maxwell et al., 1999). However, when an animal encounters hypoxic conditions, low oxygen levels impede the PHD2 catalyzed hydroxylation reaction, resulting in Hif1α stabilization, dimerization with the HLH-PAS transcription factor ARNT, and activation of target genes (Sonnenfeld et al., 1997, Lavista-Llanos et al., 2002, Bacon et al., 1998, Romero et al., 2008). As a result of this mechanism, cells are capable of coordinating and regulating cell growth and proliferation, metabolism, and development with oxygen availability.

While Hif1α is most closely associated with oxygen-sensing and hypoxia, Hif1α can also be stabilized under normoxic conditions (Magar et al., 2024, Hayashi et al., 2019). This phenomenon has become the focus of intense interest in the field of cancer metabolism, where Hif1α can be stabilized by small molecule metabolites such L-2-hydroxyglutarate (Chowdhury et al., 2011, Intlekofer et al., 2017, Nadtochiy et al., 2016), growth factor signaling (Agani and Jiang, 2013, Dekanty et al., 2005, Feldser et al., 1999), and interactions with proteins such as p53 and HSP90 (Ravi et al., 2000, Isaacs et al., 2002, Cheng et al., 2007, Liu and Semenza, 2007)(Joshi et al., 2014). The resulting pseudohypoxic state reprograms the cellular metabolism of cancer cells in a manner that promotes growth, survival, and metastasis (Hayashi et al., 2019, Magar et al., 2024).

The fly has proven to be a powerful model for studying Hif1α activity and regulation (Romero et al., 2007, Vasilia and Chrysoula, 2018). Similar to the oxygen-sensing mechanism active in mammalian cells, the *Drosophila* Hif1α homolog Similar (Sima/HIF1α) is hydroxylated by PHD2 (Fatiga) under normoxic conditions and subsequently ubiquitinated and degraded in a VHL-dependent manner (Lavista-Llanos et al., 2002, Arquier et al., 2006, Vasilia and Chrysoula, 2018). When the fly encounters a low oxygen environment, stabilized Sima/HIF1α is transported to the nucleus, where together with its heterodimeric binding partner Tango/ARNT, activates expression of target genes (Lavista-Llanos et al., 2002, Vasilia and Chrysoula, 2018, Romero et al., 2008, Li et al., 2013). Moreover, an increasing number of *Drosophila* studies have demonstrated that Sima/HIF1α can be stabilized under normoxic conditions to drive cell fate decisions. For example, studies in a wing imaginal disc tumor model revealed that activation of the oncogene PDGF/VEGF-receptor stabilizes Sima/HIF1α independent of oxygen availability, which in turn, activates a pseudo-hypoxic state, induces Ldh expression, and supports tumor growth (Wang et al., 2016). Similarly, normoxic Sima/HIF1α stabilization plays an essential role in the differentiation of blood cells within the *Drosophila* lymph gland (Mukherjee et al., 2011). Overall, these findings demonstrate that Sima/HIF1α can activate glycolytic gene expression in the fly, regulate cell fate decisions, and generally function in a manner analogous to mammalian HIF1α homologs.

While both ERR and Sima/HIF1α are well-known to regulate cellular metabolism, the interaction between these two ancient families of transcription factors remains poorly studied. This lack of focus on the interaction between ERR and Sima/HIF1α is surprising considering that previous studies in both flies and mammals demonstrated that HIF1α protein can physically bind to ERR family members (Ao et al., 2008, Zou et al., 2014), suggesting that these transcription factors cooperatively regulate gene expression. Moreover, the interaction between human ERR family members and Hif1α has been demonstrated to be functionally significant, as ERRα and Hif1α stabilize one another and synergistically promote tumor growth in a xenograft model (Ao et al., 2008). The *in vivo* significance of this ERR-Hif1α interaction, however, remains largely unexplored in both flies and mammals. The reason for a lack of follow-up studies in *Drosophila* stems, in part, from a previous study which demonstrated Sima/HIF1α and dERR regulate distinct transcriptional and metabolic programs during late L3 larval development (Li et al., 2013). Moreover, unlike *dERR* mutants, which die as mid-L2 larvae, *sima* mutants exhibit a milder growth phenotype and successfully complete larval development (Centanin et al., 2005). Together, these phenotypes suggest that Sima/HIF1α and dERR regulate independent metabolic programs during larval development.

For nearly two decades, the most commonly used *sima* allele has been *sima^KG07607^*, which has been repeatedly used for studies of growth, development, and metabolism (Centanin et al., 2005, Mortimer and Moberg, 2009, Mortimer and Moberg, 2013, Li et al., 2013, Bandarra et al., 2015, Vigne and Frelin, 2008, Lee et al., 2019, Bailey et al., 2015, Tamamouna et al., 2021, Texada et al., 2019). *sima^KG07607^*, however, is likely a hypomorphic allele, as the causal mutation consists of a p-element insertion in the first *sima* intron (Centanin et al., 2005, Öztürk-Çolak et al., 2024). Since two of the four Sima/HIF1α isoforms are composed of exons located downstream of *sima^KG07607^*, we hypothesized that Sima/HIF1α has molecular functions that were overlooked in earlier studies. We therefore used CRISPR/Cas9 to generate more severe loss-of-function mutations within the *sima* coding region and analyzed the resulting mutants for developmental phenotypes. Our analysis revealed that larvae harboring these novel *sima* alleles phenocopy the metabolic and developmental defects displayed by *dERR* mutants. Specifically, we find that *sima* mutants fail to activate aerobic glycolysis during the latter half of embryogenesis, and as a result, die during the L2 stage with severe defects in carbohydrate metabolism. Considering that that HIF1α and ERR proteins can physically interact, our findings raise the possibility that Sima/HIF1α and dERR cooperatively regulate the larval glycolytic program. Consistent with this possibility, we find that loss of either dERR or Sima/HIF1α results in diminished expression of the other protein within the larval brain. Overall, our studies indicate that both Sima/HIF1α and dERR promote juvenile growth by establishing the larval glycolytic program and demonstrate that *Drosophila* can serve as a powerful genetic model to understand how mammalian HIF1α and ERR cooperatively activate the Warburg effect.

## METHODS

### *Drosophila* husbandry and genetics

*Drosophila* stocks were maintained on standard Bloomington *Drosophila* Stock Center (BDSC) media at 25 °C. Fly strains containing *sima^KG07607^* (RRID:BDSC_14640) and *Df(3R)BSC502* (RRID:BDSC_25006) were obtained from the BDSC. Prior to use in this study, the balancer chromosomes in both strains were replaced with *TM3, P{Dfd-GMR-nvYFP}3, Sb^1^* (from RRID:BDSC_23231). Similarly, the *sima^1^* and *sima^20^*alleles were balanced over *TM3, P{Dfd-GMR-nvYFP}3, Sb^1^*. Mutant embryos and larvae were selected based on the lack of YFP expression. The strain *w^1118^; PBac{y[+mDint2] w[+mC]=ERR-GFP.FSTF}VK00037* (RRID:BDSC_38638) was used to examine dERR expression and the strain *w^1118^; PBac{y[+mDint2] w[+mC]=sima-GFP.B.FPTB}VK00037* (RRID:BDSC_43957) was used to examine expression of the Sima/HIF1α B-isoform. These strains harbor an integrated bacterial artificial chromosome that contains either the *dERR* or *sima* genomic locus with a GFP-StrepII-FLAG tag inserted at the C-terminus (Venken et al., 2009). Strains containing the *dERR^1^* (RRID:BDSC_83688) and *dERR^2^* (RRID:BDSC_83689) alleles used in this study were previously described and are available from the BDSC.

Embryos and larvae were synchronized and collected as previously described (Li and Tennessen, 2017). Briefly, synchronized populations of embryos were collected for 4 hrs using a molasses agar plate with yeast paste smeared on the surface. Embryos were allowed to hatch on the surface of the collection plate and larvae were reared on the same plate in a 25 °C incubator.

Flybase was used as an essential reference tool throughout this study (Gramates et al., 2022, Öztürk-Çolak et al., 2024).

### Generation of *sima* mutations

*sima* mutations were generated using CRISPR/Cas9. A guide RNA construct that targeted a region of *sima* (5′-GAGATCCGTCTCCGTTAGCAGGG-3′) was designed using the Fly CRISPR Optimal Target Finder (Gratz et al., 2014). Oligonucleotides containing these sequences were inserted into pU6-BbsI-chiRNA (RRID:Addgene_45946), and the resulting plasmid was injected into BDSC Stock #52669 (RRID:BDSC_52669) by Rainbow Transgenic Flies. Mutations were identified using a PCR-based method and two mutations were recovered, *sima^1^* with 1 bp deletion and *sima^20^* with a 20 bp deletion.

### Metabolic Assays

Trehalose, triglyceride, and ATP assays were conducted as previously described. (Tennessen et al., 2014a). For all assays, 25 L-2 were collected and washed with ice-cold PBS (3 times). For the ATP assay, larvae were homogenized in 100 μl of homogenization buffer [6 M guanidine HCl, 100 mM Tris (pH 7.8), 4 mM EDTA], and ATP measurement was performed using ATP measurement kit (Molecular Probes ATP kit; A22066). NAD/NADH levels were measured as previously reported (Li et al., 2017).

### Sample Preparation for Metabolomic Analysis

Larvae were collected for metabolomic studies as previously described (Li and Tennessen, 2018). Briefly, 25 L2 larvae were collected in 1.5 mL microfuge tubes and washed with twice with ice-cold PBS and once with ice-cold water. Samples tubes were immediately flash-frozen in liquid nitrogen and stored at −80°C until processing. Prior to extraction, larval pellets were transferred to a pre-tared 2 mL screwcap tube containing 1.4 mm ceramic beads. The sample mass was recorded, and the tubes immediately placed back into liquid nitrogen. Tubes were then transferred to a -20°C tube caddy and 800 µl of prechilled 90% methanol containing 1.25 µg/ml succinic-d4acid (Sigma-Aldrich; 293075) was added to each tube. Samples were immediately homogenized using an Omni BeadRuptor 24. The homogenized sample was placed at -20°C for one hour, centrifuged at 20,000xg for 5 minutes, and the supernatant was transferred to new 1.5 mL microfuge tube. The supernatant was dried in a vacuum centrifuge overnight and shipped on dry ice to the University of Colorado School of Medicine for metabolomics analysis.

### Ultra High-pressure Liquid Chromatography - Mass Spectrometry (UHPLC-MS)-based Metabolomics

UHPLC-MS metabolomics analyses were performed at the University of Colorado Anschutz Medical Campus. Briefly, the analytical platform employs a Vanquish UHPLC system (Thermo Fisher Scientific, San Jose, CA, USA) coupled online to a Q Exactive mass spectrometer (Thermo Fisher Scientific, San Jose, CA, USA). The (semi)polar extracts were resolved over a Kinetex C18 column, 2.1 x 150 mm, 1.7 µm particle size (Phenomenex, Torrance, CA, USA) equipped with a guard column (SecurityGuard^TM^ Ultracartridge – UHPLC C18 for 2.1 mm ID Columns – AJO-8782 – Phenomenex, Torrance, CA, USA) using an aqueous phase (A) of water and 0.1% formic acid and a mobile phase (B) of acetonitrile and 0.1% formic acid for positive ion polarity mode, and an aqueous phase (A) of water:acetonitrile (95:5) with 1 mM ammonium acetate and a mobile phase (B) of acetonitrile:water (95:5) with 1 mM ammonium acetate for negative ion polarity mode. The Q Exactive mass spectrometer (Thermo Fisher Scientific, San Jose, CA, USA) was operated independently in positive or negative ion mode, scanning in Full MS mode (2 μscans) from 60 to 900 m/z at 70,000 resolution, with 4 kV spray voltage, 45 sheath gas, 15 auxiliary gas. Calibration was performed prior to analysis using the Pierce^TM^ Positive and Negative Ion Calibration Solutions (Thermo Fisher Scientific).

### Statistical Analysis of Metabolomics Data

Both metabolomics datasets were analyzed using Metaboanalyst 6.0 (Pang et al., 2024), with data normalized to sample mass and preprocessed using log normalization and Pareto scaling.

### RNAseq Analysis

For larval RNA-seq analysis, 25 L2 larvae were collected and washed twice with ice-cold PBS and once with ice-cold water. For embryonic RNA seq analysis, 200 embryos were collected and washed with ice-cold PBST(twice) and ice-cold water (once). Samples were homogenized, and total RNA was extracted using the RNeasy kit (Qiagen; 74104). Sequencing and data analysis was conducted in the Indiana University Center for Genomics and Bioformatics.

Analysis was performed as described previously using a python based in-house pipeline (https://github.com/jkkbuddika/RNA-Seq-Data-Analyzer) (Buddika et al., 2021). First, the quality of raw sequencing files was assessed using FastQC version 0.11.9 (Andrews, 2010), and reads with low quality were removed using Cutadapt version 2.9 (Martin, 2011). Subsequently, the remaining high-quality reads were aligned to the Berkeley *Drosophila* Genome Project (BDGP) assembly release 6.32 (Ensembl release 103) reference genome using STAR genome aligner version 2.7.3a (Dobin et al., 2013). Additionally, duplicated reads were eliminated using SAMtools (Li et al., 2009) version 1.10. Finally, the Subread version 2.0.0 (Liao et al., 2019), function *featureCounts*, was used to count the number of aligned reads to the nearest overlapping feature. All subsequent downstream analyses and data visualization steps were performed using custom scripts written in R. To identify differentially expressed genes in different genetic backgrounds the Bioconductor package DESeq2 (https://bioconductor.org/packages/release/bioc/html/DESeq2.html) version 1.26.0 was used (Love et al., 2014). Unless otherwise noted, significantly upregulated and downregulated genes were defined as FDR < 0.05; Log_2_ fold change > 1 and FDR < 0.05; Log_2_ fold change < -1.

### PANGEA Analysis

All RNAseq data were analyzed using PANGEA (Hu et al., 2023). Genes that were significantly down- or up-regulated (Log_2_ fold change>|1.0|, adj *P-*value<0.005) were analyzed for Gene Ontology Enrichment using the GO Biological Processes and REACTOME pathway sets (Consortium et al., 2023).

### Immunofluorescence

Larvae were dissected in 1x phosphate buffered saline (PBS) and fixed for 30 minutes in 4% paraformaldehyde in 1x PBS at room temperature. Fixed tissues were then washed 3 times for 5 minutes each with PT (PBS with 0.1% Triton X-100) and blocked for 30 minutes in NGS-PT (PT with 5% normal goat serum (NGS)) at room temperature. Blocked tissues were incubated overnight at 4 °C with rabbit anti-GFP (#A11122 Thermo Fisher) diluted 1:500 in NGS-PT. Tissues were subsequently washed 3 times for 5 minutes each with PT and then incubated overnight with Alexa Fluor 568 donkey anti-rabbit secondary antibody (#A10042 Thermo Fisher) diluted 1:1000 in PT at 4 °C. Tissues were then washed 3 times for 5 minutes each in PT and mounted in Vectashield with DAPI (Vector Laboratories; H-1200-314 10).

### Data Availability

All strains and reagents are available upon request. All targeted metabolomics data described herein are included in Table S1. Processed RNA-seq data is presented in Tables S5-S7 and available in NCBI Gene Expression Omnibus (GEO; GSE284097).

## RESULTS

### Sima is essential for *Drosophila* larval development

*Drosophila* Sima/HIF1α has been extensively studied in the context of hypoxia signaling, metabolism, and development (Centanin et al., 2005, Vasilia and Chrysoula, 2018, Romero et al., 2007). These studies, however, largely rely on the *sima^KG07607^*allele, which represents a p-element insertion in one of the upstream *sima* introns (Figure 1A)(Centanin et al., 2005). Since at least one *sima* isoform is produced from coding regions located downstream of this p-element insertion, *sima^KG07607^* is likely not a null allele, thus raising the possibility that certain roles for Sima/HIF1α in regulating larval development remain undiscovered. We examined this possibility using CRISPR/Cas9 to generate two new *sima* alleles that induce small deletions and frameshifts within the coding region (Figure 1A; see Methods).

**Figure 1.**
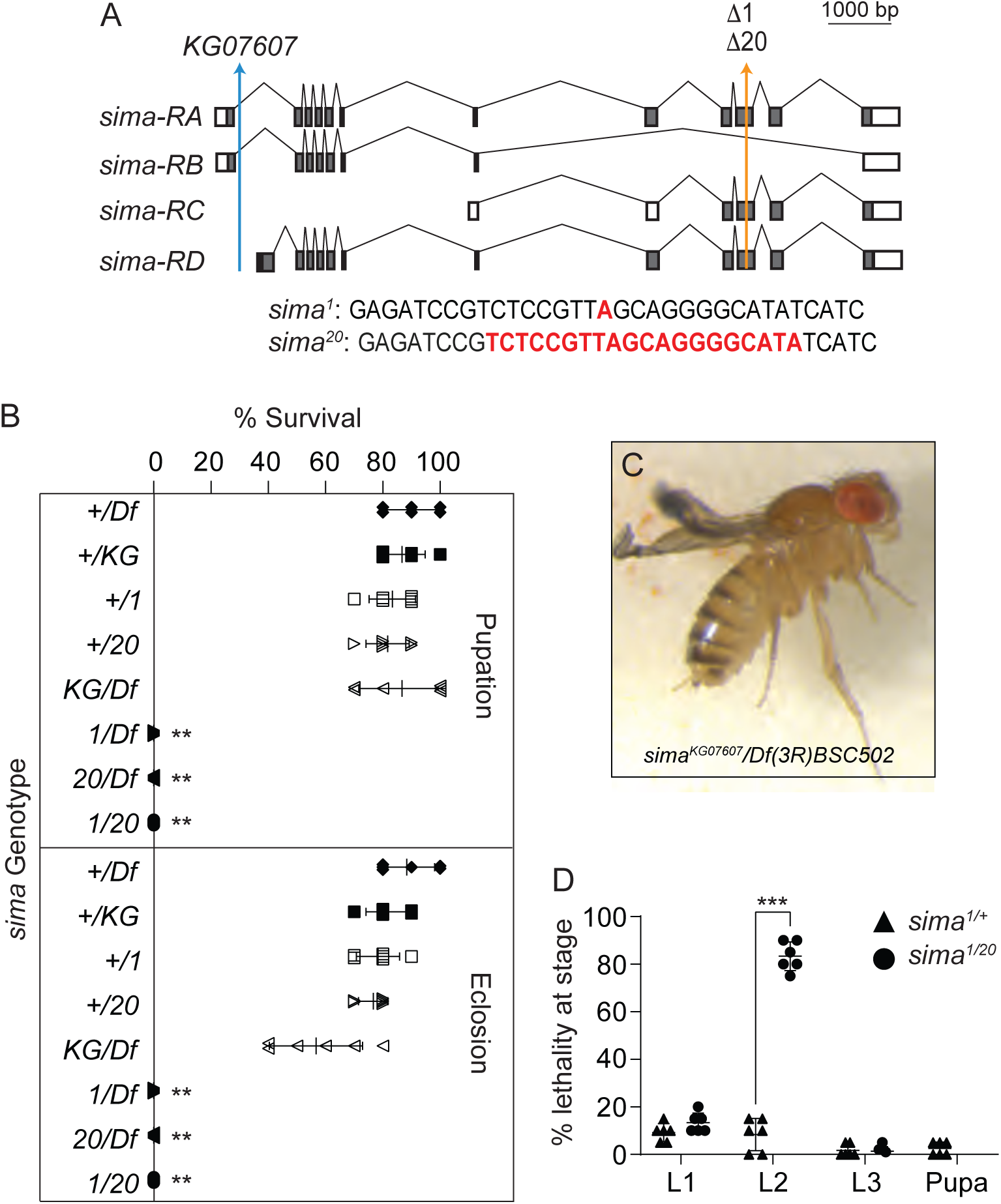
Loss-of-function mutations within the *sima* coding region induce a larval lethal phenotype. (A) A schematic diagram illustrating the *sima* locus and the mutations *sima^KG07607^*, *sima^Δ1^* (1 bp deletion), and *sima^Δ20^*(20 bp deletion). (B) The percentage of animals that completed larval development and metamorphosis were quantified for the indicated genotypes. + indicates a wild-type allele from *w^1118^*. The remaining alleles are abbreviated as indicated: *Df(3R)BSC502* (Df), *sima^KG07607^* (KG) *sima^Δ1^* (1) or *sima^Δ20^* (20). (C) *sima^KG07607^/Df(3R)BSC502* adults exhibit a wrinkled wing phenotype. (D) Lethal phase analysis of *sima* mutants. *sima^Δ1/+^* controls and *sima^Δ1/20^*mutants were collected as newly hatched first instar larvae and followed through development, scoring for the percentage of animals that die as either first instar (L1), second instar (L2), or third instar (L3) larvae. For (B) and (D), n=6 vials containing 20 embryos for each genotype. \*\**P*<0.001. For (B), *P* value was calculated relative to the *Df/+* genotype using a Kruskal-Wallis test followed by a Dunn’s multiple comparison test. For (D), *P* value was calculated using an unpaired student’s t-test.

To determine if the new *sima* alleles induced previously undescribed developmental defects, we independently crossed *sima^1^*, *sima^20^*, and *sima^KG07607^* with the flies containing the deficiency *Df(3R)BSC502*, which removes the entire *sima* locus. Consistent with earlier studies, *sima^KG07607^/Df* larvae were viable and successfully completed both larval and pupal development (Figure 1B), with adult mutant flies exhibiting a previously described wing defect (Figure 1C)(Centanin et al., 2005). In contrast, trans-heterozygous *sima^1^/Df*, *sima^20^/Df*, and *sima^1/20^* animals died during larval development (Figure 1B). Moreover, a viability analysis of staged *sima^1^*^/*20*^ mutants revealed that these new *sima* alleles induce a L2 lethal phase (Figure 1D). Overall, our results demonstrate that Sima/HIF1α is required for normal larval development and confirm that *sima^KG07607^*is a hypomorphic allele.

### *sima* mutant larvae exhibit defects in carbohydrate metabolism

The *sima* mutant mid-L2 lethal phenotype is reminiscent of *dERR* mutants, where a failure to increase glycolytic metabolism at the onset of larval development renders the larvae unable to use carbohydrates for growth and energy production (Tennessen et al., 2011). Since dERR and Sima/HIF1α proteins are also known to physically interact (Li et al., 2013), we examined the possibility that *sima* mutants also exhibit a disruption of the larval glycolytic program. As a first step towards analyzing this possibility, we measured levels of the circulating disaccharide trehalose in both *sima^1/20^* mutants and *sima^1/+^* heterozygous control larvae. Consistent with Sima/HIF1α regulating larval carbohydrate metabolism, we found that trehalose levels of mid-L2 *sima* mutants were elevated more than 2-fold when compared to controls (Figure 2A). Moreover, although the NAD/NADH ratio was normal in *sima* mutants (Figure 2B), ATP levels were significantly decreased (Figure 2C), suggesting that *sima* mutant larvae are unable to properly utilize sugars for energy production (Figure 2C). We would note, however, that unlike *dERR* mutants, *sima* mutants exhibited normal TAG levels, indicating that *sima* larvae are still able to maintain normal lipid storage under normoxic conditions (Figure 2D).

**Figure 2.**
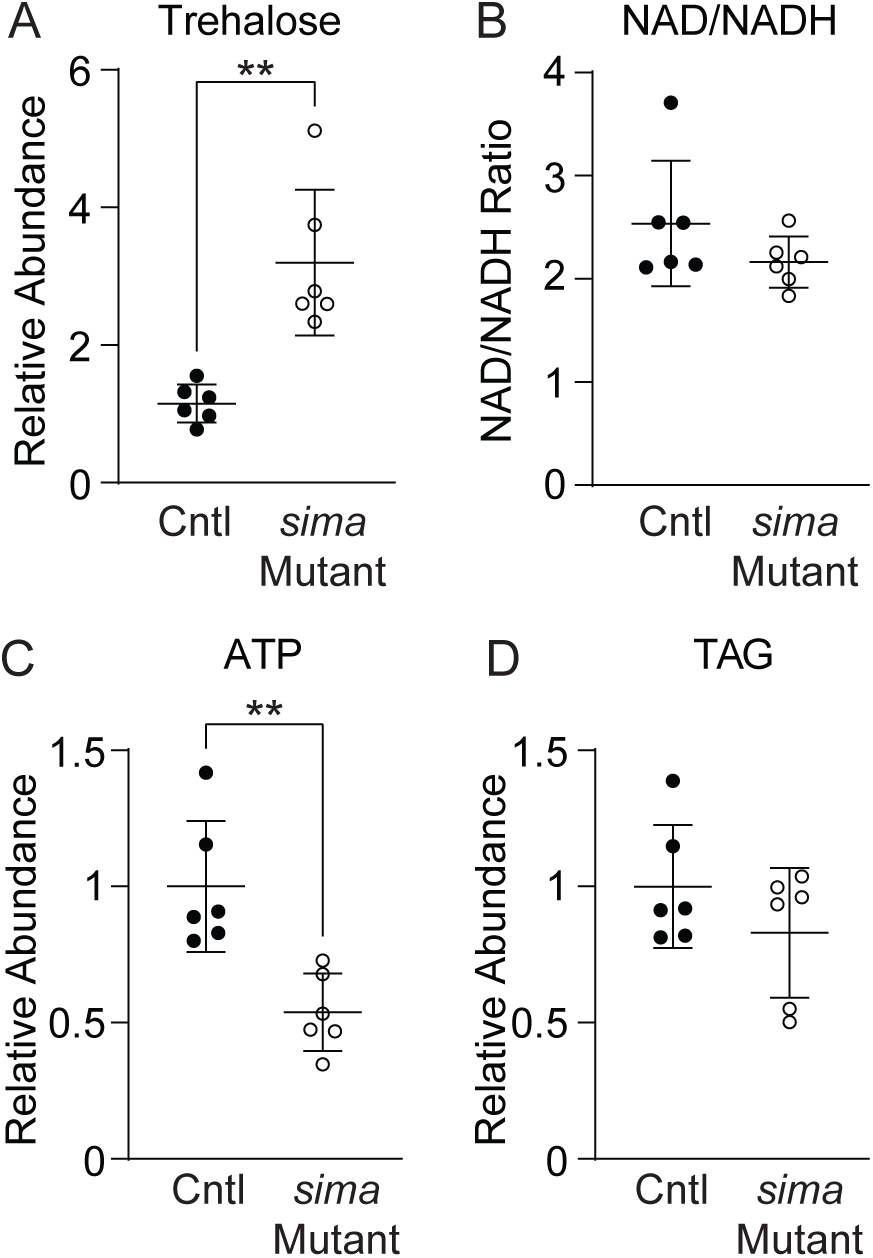
*sima* mutant larvae accumulate excess trehalose and harbor decreased ATP levels. When compared with *sima^1/+^* heterozygous controls, *sima^1/20^* mutant larvae display (A) a significant increase in trehalose level, (B) a significant decrease in ATP, and (C,D) no significant differences in either the NAD/NADH ratio or TAG abundance. Data are normalized to soluble protein. n=6 biological replicates containing 25 mid-L2 larvae per sample. \*\**P*<0.01 using an unpaired student’s t-test.

To further explore the *sima* mutant metabolic phenotypes, we used a semi-targeted metabolomics approach to compare *sima^1^*^/*20*^ mutants with *sima^1^*^/+^ heterozygotes at the mid-L2 larval stage (Table S1). Our analysis revealed a significant increase in trehalose and several other carbohydrates (Figure 3A-D), thus confirming our enzyme-based trehalose measurements (Figure 2A) and again suggesting that *sima* mutants experience defects in carbohydrate metabolism. Moreover, lactate and 2-hydroxyglutarate (2HG; Figure 3A,B,E,F) were significantly decreased in *sima* mutants. This result is of particular significance because these two metabolites are also significantly altered in *dERR* mutants (Li et al., 2017, Tennessen et al., 2011). Overall, our findings not only revealed that *sima* mutants harbor defects in glycolytic metabolism but also demonstrated that the *sima* mutant metabolomic profile mirrors that of *dERR* mutant larvae.

**Figure 3.**
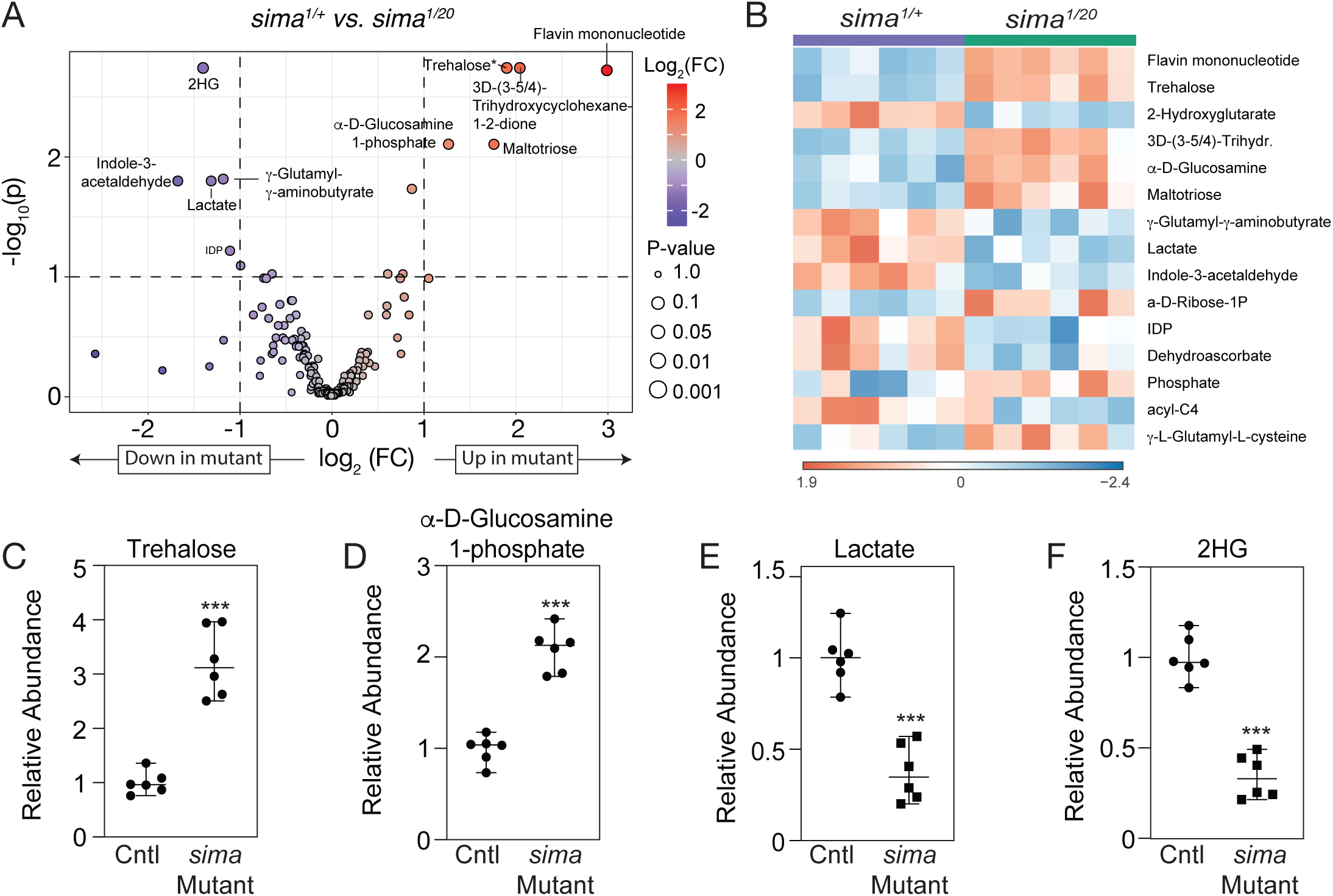
Metabolomic analysis of *sima* mutant larvae. *sima^1/20^* mutant larvae and *sima^1/+^* controls were analyzed using a semi-targeted LC-MS based method. (A) Differences in metabolite abundance between control and mutant samples are represented as a volcano plot. Dashed vertical line represents a log_2_ fold change (FC) of > 2. Dashed horizontal line represents p < 0.1. (B) A heat map illustrating the top 15 most significantly altered compounds in *sima^1/20^* mutants as compared with *sima^1/+^*controls. Panels (A) and (B) were generated using Metaboanalyst 6.0, as described in the methods. (C-F) Boxplots illustrating the normalized abundance of metabolites that are significantly altered in *sima^1/20^* mutant larvae relative to *sima^1/+^* controls. Black dots represent individual samples and the horizontal bar in the middle represents the median. 2-hydroxyglutarate is abbreviated 2HG. n=6 biological replicates containing 25 mid-L2 larvae per sample. \*\**P*<0.001 using an unpaired student’s t-test.

### *Drosophila* Sima/HIF1α is an essential regulator of glycolysis during normoxic larval development

The metabolomics analysis described above indicates the *sima* mutants are unable to properly metabolize carbohydrates. To further explore the metabolic basis of this phenotype, we used RNAseq to compare gene expression between *sima* mutants and heterozygous controls in mid-L2 larvae. A total of 781 genes were misregulated >2-fold in *sima* mutants (p < 0.005; Log_2_ fold change > 1 and p < 0.005; Log_2_ fold change < -1; Figure 4A; Table S2, S3). Included among the significantly downregulated genes was nearly every gene directly involved in glycolysis, including genes that encode the rate-limiting enzymes Hexokinase A (HexA), Phosphofructokinase (Pfk), and Pyruvate kinase (Pyk) (Figure 4B). In addition, *Lactate dehydrogenase* (*Ldh*, also known as *ImpL3*) was significantly down-regulated in our dataset (Log_2_-fold change of -3.6; Table S2). Considering that Ldh catalyzes the formation of both L-lactate and L-2-hydroxyglutarate (Li et al., 2017), the decrease in *Ldh* gene expression is consistent with the decreased abundance of both metabolites in the metabolomics data (see Figure 3).

**Figure 4.**
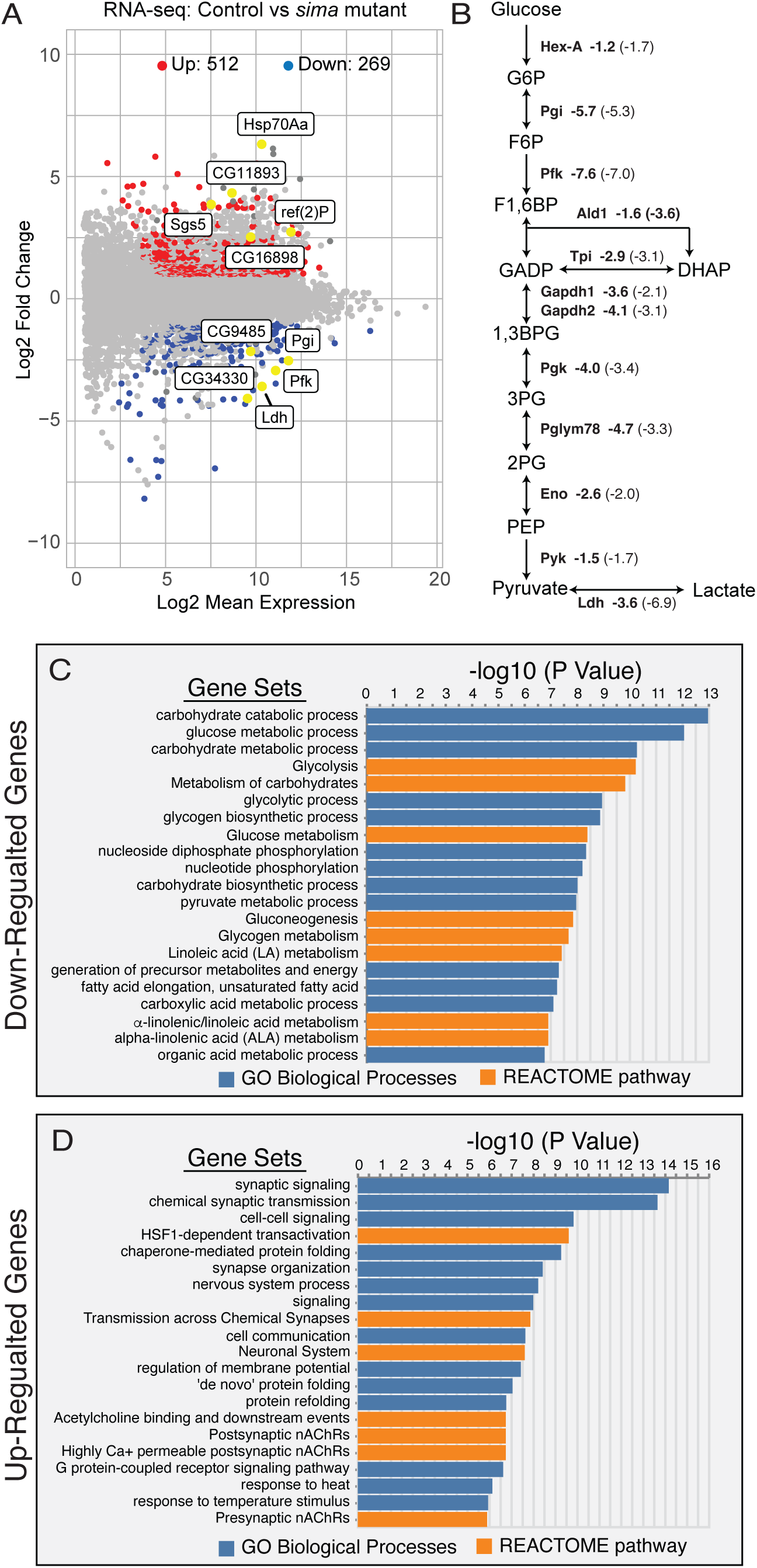
Expression of genes that encode glycolytic enzymes are significantly downregulated in *sima* mutants. (A) A volcano plot illustrating changes in gene expression in *sima^Δ1/20^*mutant larvae relative to *sima^Δ1^/+* controls. (B) A diagram of glycolysis is depicted that displays the glycolytic genes that are downregulated in *sima* mutants followed by their log2 fold change in expression (in bold) compared to what reported in ERR mutants (in parentheses). (C,D) PANGEA was used to determine GO enrichment of biological processes among those genes that were either (C) down-regulated or (D) up-regulated in *sima^Δ1/20^* mutants.

As an extension of our targeted analysis of glycolytic genes, we also used PAthway, Network, and Gene-set Enrichment Analysis (PANGEA) to conduct a Gene Ontology (GO) analysis of the *sima* mutant RNA-seq dataset. These studies revealed a significant enriched for genes involved in carbohydrate metabolism – in fact, of the top 20 most significantly enriched GO terms among the down-regulated genes, 12 are associated with carbohydrate metabolism or a related process (Figure 4C). In contrast, PANGEA analysis of significantly up-regulated genes failed to identify significant enrichment for metabolism-related GO categories. Instead, the analysis revealed enrichment for genes associated with neuronal function (e.g., synaptic signaling) and the stress response, such as the heat-shock proteins that fall under the GO term “HSF1-dependent transactivation” (Figure 4D, Table S4).

Overall, the general gene expression changes measured in *sima* mutants are reminiscent of those observed in *dERR* mutants. In fact, the decrease in glycolytic genes is nearly identical in *dERR* and *sima* mutants, with nearly 50% of the genes that are down-regulated in *sima* mutants being previously reported as down-regulated in *dERR* mutants (Figure 4B, Figure S1; Tennessen et al., 2011). For example, *Pfk* was down-regulated -7.0-fold in *dERR* mutants and -7.6-fold in *sima* mutants (Figure 4B). These observations support a model in which dERR and Sima/Hif1α cooperatively regulate larval gene expression.

### *sima* mutants fail to activate aerobic glycolysis during the embryonic metabolic transition

The larval glycolytic program stems from a metabolic switch that occurs approximately midway through embryogenesis (i.e., the embryonic metabolic transition; EmbMT) (Tennessen et al., 2014b), when dERR coordinately activates the expression of glycolysis and related pathways in carbohydrate metabolism. Considering the similarities between the *sima* and *dERR* mutant phenotypes, we next examined the possibility that Sima/Hif1α is also required to activate the EmbMT. We tested this hypothesis by collecting *sima* mutant embryos and heterozygous control embryos at 4-hour intervals, from the onset of the EmbMT (8-12 hr after egg-laying [AEL]) until near the end of embryogenesis (16-20 hrs AEL). If Sima/Hif1α were required during this developmental event, expression of the genes that encode glycolytic enzymes should fail to be upregulated in the 8-12 hr samples. In contrast, if Sima/Hif1α were simply required to maintain glycolytic gene expression after activation of the EmbMT, we would expect to see that those genes encoding glycolytic enzymes would initially increase in expression but then decrease during larval development. Our studies support the first model, as RNA-seq analysis revealed that nearly every glycolytic gene that is transcriptionally up-regulated at 8-12 hrs is expressed at a lower level in *sima* mutants relative to controls at all three timepoints (Figure 5 and Table S5,S6,S7,S8,S9,S10). Moreover, of those glycolytic genes that increased in expression in *sima* mutant embryos, this increase was significantly lower than in the control group (Figure 5 and Table S5, S6, S7, S8, S9, S10).

**Figure 5.**
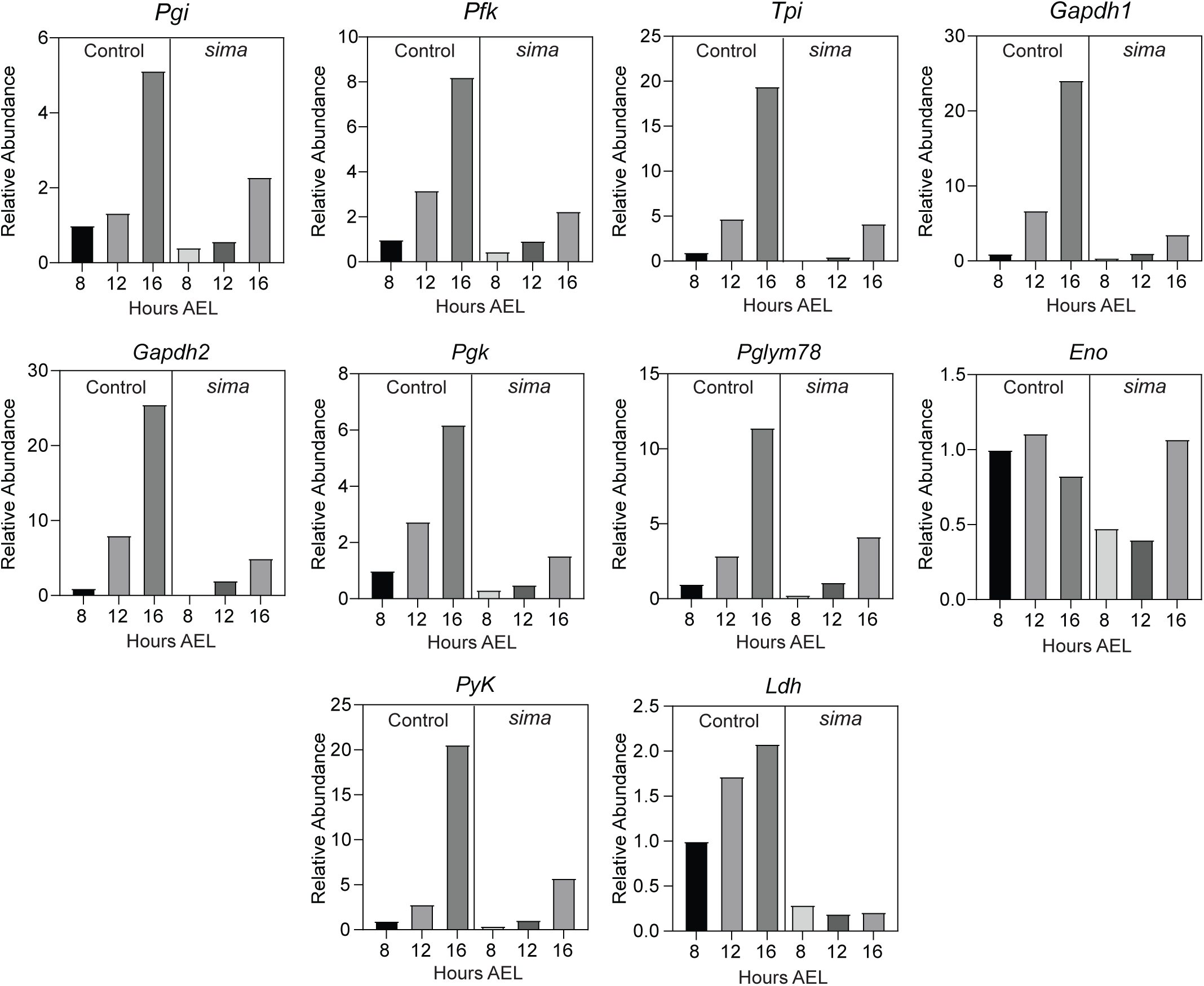
*Sima* mutants fail to activate aerobic glycolysis during the embryonic metabolic transition. RNA was collected from synchronized populations of *Sima^Δ1/+^* controls and *Sima^Δ1/20^* mutants at 8-12, 12-16, and 16-20 hours after egg laying (AEL) and subsequently analyzed by RNAseq. Expression data from each gene was normalized to control group at 8-12 hours AEL. *sima* mutants exhibit a significant reduction in glycolytic gene expression at all time points. n=3 biological replicates containing 200 embryos per sample. See Tables S5, S6, and S7 for significance values of these changes.

Our RNAseq data indicate that Sima/Hif1α plays a critical role in the activation of aerobic glycolysis in preparation for larval development. To further explore the nature of the Sima/Hif1α embryonic transcriptional program, we analyzed the embryonic RNA-seq data from *sima* mutants using PANGEA. When compared with the *sima^1/+^* heterozygous control, genes that were significantly down-regulated in *sima^1/20^* mutants displayed enrichment for GO categories related to glycolytic metabolism at all three embryonic timepoints (Table 1, S11, S12, S13). In fact, among the top 10 most significantly enriched gene set categories, nearly all correspond to glycolysis, carbohydrate metabolism, or a related metabolic pathway. The one notable exception to this trend is in the 8-12 hr embryonic dataset, where genes involved in the heat-shock response are also significantly down-regulated. This result is of considerable interest, as a previous report indicated that Sima/Hif1α was required for normal activity of the *heat shock factor* (*hsf*) transcription factor (Baird et al., 2006). In contrast, the gene set categories that are significantly enriched across the significantly up-regulated genes varies with by timepoint and are generally unrelated to metabolism, although there is a general trend for enrichment of terms related to cuticle development and the cell cycle (Table 2, S11, S12, S13).

**Table 1.** PANGEA analysis of significantly down-regulated genes in *sima[1/20]* mutants as compared with *sima[1/+]* heterozygous controls. The top 10 most significantly enriched Gene Set Categories are listed for each timepoint.

| PANGEA analysis of genes down-regulated in *sima* mutants (8-12 hrs after egg laying) |  |  |  |  |  |  |  |
| --- | --- | --- | --- | --- | --- | --- | --- |
| Gene Set Category | Gene Set ID | Gene Set Name | Gene Set Size | % Genes in Gene Set | Fold Enrichment | log2 fold | P value |
| REACTOME pathway | R-DME-3371571 | HSF1-dependent transactivation | 30 | 8.8 | 40.8 | 5.4 | 2.3E-14 |
| GO Biological Processes | GO:0006096 | glycolytic process | 27 | 7.9 | 40.8 | 5.4 | 4.8E-13 |
| REACTOME pathway | R-DME-70171 | Glycolysis | 28 | 7.9 | 39.4 | 5.3 | 7.1E-13 |
| GO Biological Processes | GO:0006165 | nucleoside diphosphate phosphorylation | 31 | 7.9 | 35.6 | 5.2 | 2.0E-12 |
| GO Biological Processes | GO:0046939 | nucleotide phosphorylation | 32 | 7.9 | 34.5 | 5.1 | 2.8E-12 |
| GO Biological Processes | GO:0016052 | carbohydrate catabolic process | 56 | 8.8 | 21.9 | 4.5 | 2.3E-11 |
| REACTOME pathway | R-DME-70326 | Glucose metabolism | 41 | 7.9 | 26.9 | 4.7 | 3.3E-11 |
| REACTOME pathway | R-DME-3371556 | Cellular response to heat stress | 79 | 9.6 | 17.1 | 4.1 | 3.7E-11 |
| GO Biological Processes | GO:0006090 | pyruvate metabolic process | 45 | 7.9 | 24.5 | 4.6 | 8.1E-11 |
| REACTOME pathway | R-DME-70263 | Gluconeogenesis | 25 | 6.1 | 34.3 | 5.1 | 8.5E-10 |
| PANGEA analysis of genes down-regulated in *sima* mutants (12-16 hrs after egg laying) |  |  |  |  |  |  |  |
| Gene Set Category | Gene Set ID | Gene Set Name | Gene Set Size | % Genes in Gene Set | Fold Enrichment | log2 fold | P value |
| GO Biological Processes | GO:0006096 | glycolytic process | 27 | 8.8 | 45.6 | 5.5 | 1.7E-13 |
| REACTOME pathway | R-DME-70171 | Glycolysis | 28 | 8.8 | 44.0 | 5.5 | 2.5E-13 |
| GO Biological Processes | GO:0006165 | nucleoside diphosphate phosphorylation | 31 | 8.8 | 39.8 | 5.3 | 7.3E-13 |
| GO Biological Processes | GO:0046939 | nucleotide phosphorylation | 32 | 8.8 | 38.5 | 5.3 | 1.0E-12 |
| GO Biological Processes | GO:0016052 | carbohydrate catabolic process | 56 | 9.8 | 24.5 | 4.6 | 7.4E-12 |
| REACTOME pathway | R-DME-70326 | Glucose metabolism | 41 | 8.8 | 30.1 | 4.9 | 1.2E-11 |
| GO Biological Processes | GO:0006090 | pyruvate metabolic process | 45 | 8.8 | 27.4 | 4.8 | 2.9E-11 |
| REACTOME pathway | R-DME-71387 | Metabolism of carbohydrates | 155 | 12.7 | 11.5 | 3.5 | 9.3E-11 |
| REACTOME pathway | R-DME-70263 | Gluconeogenesis | 25 | 6.9 | 38.3 | 5.3 | 3.9E-10 |
| GO Biological Processes | GO:0005975 | carbohydrate metabolic process | 264 | 14.7 | 7.8 | 3.0 | 7.6E-10 |
| PANGEA analysis of genes down-regulated in *sima* mutants (16-20 hrs after egg laying) |  |  |  |  |  |  |  |
| Gene Set Category | Gene Set ID | Gene Set Name | Gene Set Size | % Genes in Gene Set | Fold Enrichment | log2 fold | P value |
| GO Biological Processes | GO:0016052 | carbohydrate catabolic process | 56 | 16.0 | 39.9 | 5.3 | 1.1E-16 |
| GO Biological Processes | GO:0006096 | glycolytic process | 27 | 12.0 | 62.1 | 6.0 | 9.8E-15 |
| REACTOME pathway | R-DME-70171 | Glycolysis | 28 | 12.0 | 59.9 | 5.9 | 1.4E-14 |
| GO Biological Processes | GO:0006165 | nucleoside diphosphate phosphorylation | 31 | 12.0 | 54.1 | 5.8 | 4.1E-14 |
| GO Biological Processes | GO:0046939 | nucleotide phosphorylation | 32 | 12.0 | 52.4 | 5.7 | 5.7E-14 |
| REACTOME pathway | R-DME-70326 | Glucose metabolism | 41 | 12.0 | 40.9 | 5.4 | 6.9E-13 |
| REACTOME pathway | R-DME-71387 | Metabolism of carbohydrates | 155 | 17.3 | 15.6 | 4.0 | 1.6E-12 |
| GO Biological Processes | GO:0006090 | pyruvate metabolic process | 45 | 12.0 | 37.2 | 5.2 | 1.7E-12 |
| GO Biological Processes | GO:0005975 | carbohydrate metabolic process | 264 | 20.0 | 10.6 | 3.4 | 7.8E-12 |
| REACTOME pathway | R-DME-70263 | Gluconeogenesis | 25 | 9.3 | 52.1 | 5.7 | 4.3E-11 |

**Table 2.** PANGEA analysis of significantly up-regulated genes in *sima[1/20]* mutants as compared with *sima[1/+]* heterozygous controls. The top 10 most significantly enriched Gene Set Categories are listed for each timepoint.

| PANGEA analysis of genes up-regulated in *sima* mutants<br>8-12 hrs after egg laying |  |  |  |  |  |  |  |
| --- | --- | --- | --- | --- | --- | --- | --- |
| Gene Set Category | Gene Set ID | Gene Set Name | Gene Set Size | % Genes in Gene Set | Fold Enrichment | log2 fold | P value |
| GO Biological Processes | GO:0040003 | chitin-based cuticle development | 208 | 7.3 | 4.9 | 2.3 | 4.5E-09 |
| GO Biological Processes | GO:0042335 | cuticle development | 250 | 7.7 | 4.3 | 2.1 | 2.1E-08 |
| GO Biological Processes | GO:0010529 | negative regulation of transposition | 22 | 2.6 | 16.3 | 4.0 | 1.3E-07 |
| GO Biological Processes | GO:0010528 | regulation of transposition | 24 | 2.6 | 14.9 | 3.9 | 2.6E-07 |
| GO Biological Processes | GO:0010171 | body morphogenesis | 28 | 2.6 | 12.8 | 3.7 | 8.4E-07 |
| GO Biological Processes | GO:1900153 | positive regulation of nuclear-transcribed mRNA catabolic process, deadenylation-dependent decay | 14 | 1.8 | 18.3 | 4.2 | 4.8E-06 |
| GO Biological Processes | GO:0000289 | nuclear-transcribed mRNA poly(A) tail shortening | 21 | 1.8 | 12.2 | 3.6 | 4.3E-05 |
| GO Biological Processes | GO:0034587 | piRNA metabolic process | 17 | 1.5 | 12.0 | 3.6 | 2.8E-04 |
| GO Biological Processes | GO:0000288 | nuclear-transcribed mRNA catabolic process, deadenylation-dependent decay | 33 | 1.8 | 7.8 | 3.0 | 4.2E-04 |
| REACTOME pathway | R-DME-8964038 | LDL clearance | 21 | 1.5 | 9.7 | 3.3 | 6.6E-04 |
| 12-16 hrs after egg laying |  |  |  |  |  |  |  |
| GO Biological Processes | GO:0051276 | chromosome organization | 758 | 14.5 | 2.7 | 1.4 | 3.7E-22 |
| GO Biological Processes | GO:0000278 | mitotic cell cycle | 516 | 10.3 | 2.8 | 1.5 | 4.8E-17 |
| GO Biological Processes | GO:0000280 | nuclear division | 365 | 8.3 | 3.2 | 1.7 | 6.7E-17 |
| GO Biological Processes | GO:0000819 | sister chromatid segregation | 128 | 4.7 | 5.1 | 2.4 | 1.3E-16 |
| GO Biological Processes | GO:0007059 | chromosome segregation | 232 | 6.4 | 3.8 | 1.9 | 1.6E-16 |
| GO Biological Processes | GO:0000070 | mitotic sister chromatid segregation | 112 | 4.3 | 5.4 | 2.4 | 5.3E-16 |
| GO Biological Processes | GO:0040003 | chitin-based cuticle development | 208 | 5.9 | 3.9 | 2.0 | 1.1E-15 |
| GO Biological Processes | GO:0042335 | cuticle development | 250 | 6.4 | 3.6 | 1.8 | 3.8E-15 |
| GO Biological Processes | GO:0098813 | nuclear chromosome segregation | 211 | 5.7 | 3.8 | 1.9 | 9.6E-15 |
| GO Biological Processes | GO:0051726 | regulation of cell cycle | 316 | 7.0 | 3.1 | 1.6 | 6.0E-14 |
| 16-20 hrs after egg laying |  |  |  |  |  |  |  |
| GO Biological Processes | GO:0051276 | chromosome organization | 758 | 23.0 | 4.2 | 2.1 | 9.9E-25 |
| GO Biological Processes | GO:0000278 | mitotic cell cycle | 516 | 18.9 | 5.1 | 2.4 | 3.9E-24 |
| REACTOME pathway | R-DME-69278 | Cell Cycle, Mitotic | 256 | 12.0 | 6.6 | 2.7 | 5.3E-19 |
| REACTOME pathway | R-DME-1640170 | Cell Cycle | 274 | 12.4 | 6.3 | 2.7 | 6.1E-19 |
| GO Biological Processes | GO:0000280 | nuclear division | 365 | 13.7 | 5.3 | 2.4 | 4.4E-18 |
| GO Biological Processes | GO:0000070 | mitotic sister chromatid segregation | 112 | 8.2 | 10.3 | 3.4 | 5.7E-18 |
| GO Biological Processes | GO:0051726 | regulation of cell cycle | 316 | 12.7 | 5.6 | 2.5 | 9.7E-18 |
| GO Biological Processes | GO:0000819 | sister chromatid segregation | 128 | 8.2 | 9.0 | 3.2 | 1.5E-16 |
| GO Biological Processes | GO:0007346 | regulation of mitotic cell cycle | 190 | 9.6 | 7.1 | 2.8 | 2.9E-16 |
| GO Biological Processes | GO:0007059 | chromosome segregation | 232 | 10.3 | 6.2 | 2.6 | 9.6E-16 |

### The accumulation of Sima/HIF1α and dERR proteins are mutually dependent

Considering that *Drosophila* Sima/HIF1α and dERR were previously demonstrated to physically interact (Li et al., 2013), our findings raise the question as to whether these two proteins influence the accumulation and/or stability of each other. To investigate this possibility, we used previously described recombineered BAC transgenes to express either a dERR-FLAG-STREPII-GFP (dERR-GFP) fusion protein in a *sima* mutant background or a Sima-superfolderGFP-FLAG-PreScission-TEV-BLRP (Sima-GFP) fusion protein in a *dERR* mutant background (Venken et al., 2009). While analyzing expression of these two transgenes, we found that both were robustly expressed in the optic lobes and central brain region of L2 larvae (Figure 6A-C, G-I), and thus decided to focus our studies on these regions of the larval brain. Consistent with our genetic studies which indicate that Sima/HIF1α and dERR regulate similar metabolic programs, we found that accumulation of these two proteins is mutually dependent (Figure 6). Specifically, we found that dERR-GFP was absent within the optic lobes and central brain region of *sima* mutants (Figure 6D-F), and Sima-GFP was nearly undetectable within the same regions in *dERR* mutant brain (Figure 6J-L). Overall, these findings support a model in which the abundance of dERR and Sima/HIF1α proteins are codependent.

**Figure 6.**
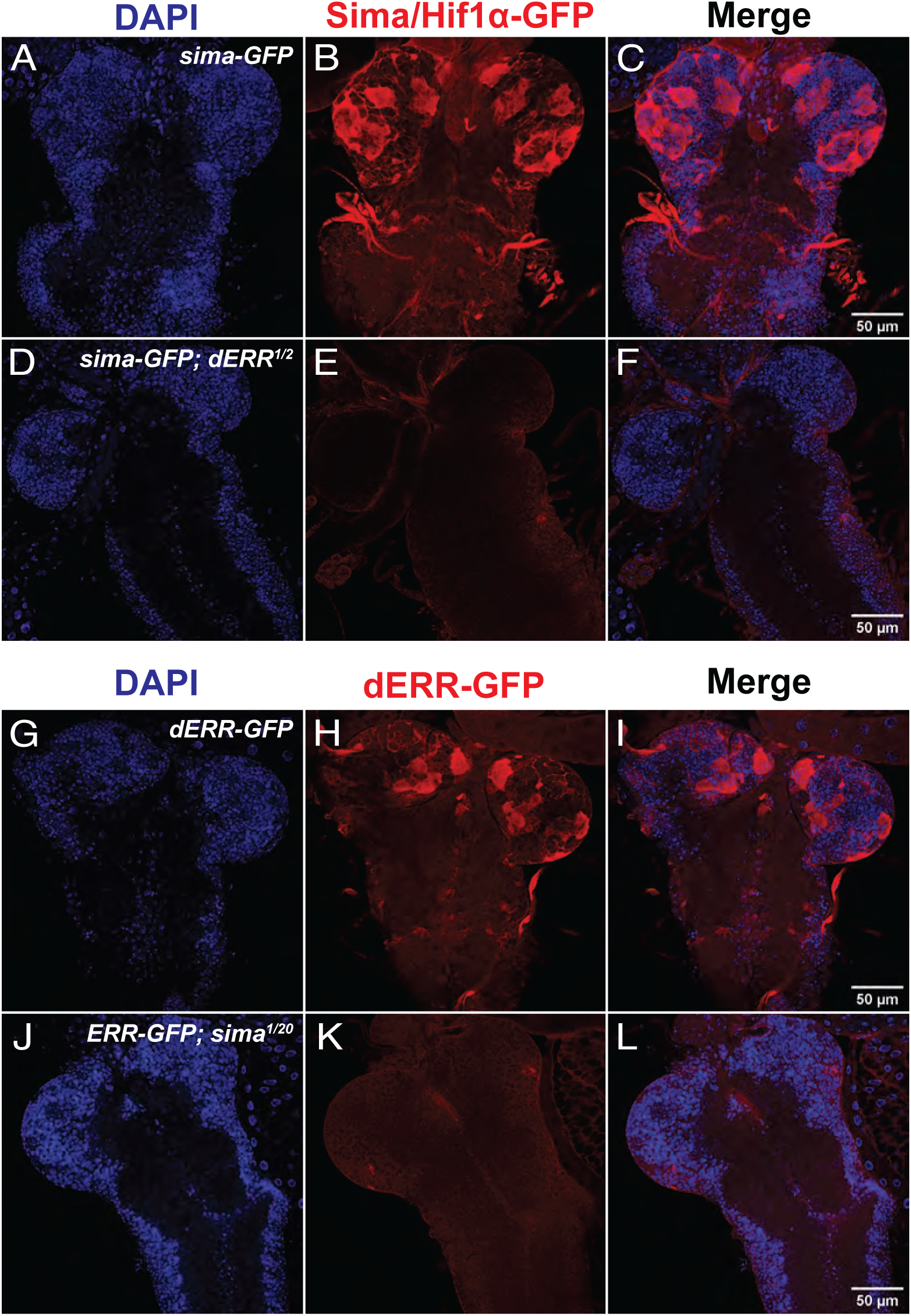
The accumulation of Sima/HIF1α and dERR proteins are mutually dependent in the larval brain. mid-L2 larvae of the indicated genotypes were collected 55 hrs after egg-laying, fixed, and stained with DAPI and an anti-GFP antibody. (A-F) The CNS from control and *dERR* mutant L2 larvae expressing Sima-GFP were stained using (A,D) DAPI and (B,E) an anti-GFP antibody. (G-L) Control and *sima* mutant L2 larvae expressing dERR-GFP were stained using (G-I) DAPI and an (J-L) anti-GFP antibody.

## DISCUSSION

Here we demonstrate that Sima/HIF1α is required during embryogenesis to coordinately up-regulate carbohydrate metabolism in preparation for larval growth. Notably, our study also reveals that the Sima-dependent gene expression profile shares considerable overlap with that observed in *dERR* mutant, suggesting that Sima/HIF1α and dERR cooperatively regulate embryonic and larval glycolytic gene expression. This finding, combined with our observation that each protein is required for accumulation of the other in the larval brain, lends credence to earlier biochemical studies which demonstrate that Sima/HIF1α and dERR physically interact (Li et al., 2013). While this earlier study was somewhat ignored due to the reported phenotypic differences between *dERR* and *sima* mutants, our study indicates that this interaction is important for the function of both transcription factors during *Drosophila* larval development.

Our findings are important because they suggest that Sima/HIF1α serve a broader role in normal *Drosophila* development than previously appreciated. While Sima/HIF1α has been intensely studied in the context of hypoxia (Lavista-Llanos et al., 2002, Centanin et al., 2005, Centanin et al., 2008, Romero et al., 2008), accumulating evidence hints at multiple functions for Sima/HIF1α in *Drosophila* development. For example, some of the earliest studies of Sima/HIF1α in the fly noted that insulin-PI3K/Tor signaling could stabilize Sima/HIF1α under normoxic conditions (Dekanty et al., 2005), thus demonstrating that specific developmental signals could potentially induce Sima/HIF1a during normoxic development. Similarly, Sima/HIF1α was subsequently found to both control the rate of border cell migration within the ovary and promote normal blood cell development within the lymph gland (Mukherjee et al., 2011, Doronkin et al., 2010). Our findings expand upon these earlier studies and suggests that Sima/HIF1α broadly regulates larval metabolism.

When considered in light of known roles for Sima/HIF1α in *Drosophila* development, our findings raise the question as to how the interaction between dERR and Sima/HIF1α activate the larval glycolytic program. Based on our observations, as well as those of previous studies (Li et al., 2013), an appealing model emerges in which these two transcription factors form a stable heterodimer and cooperatively activate gene expression. Such a model is also consistent with previous findings that Hif1α and ERRs interact in human cells. All three members of ERR family (ERRα, ERRβ and ERRγ) have been shown to interact with Hif1α, and together, this heterodimer upregulates transcriptional activation of angiogenic and glycolytic genes (Ao et al., 2008, Zou et al., 2014, Lee et al., 2012). Considering how this HIF1α-ERR interaction appears to be conserved between *Drosophila* and humans, the fly represents a promising model to better understand how these two conserved transcription factors cooperatively regulate aerobic glycolysis.

We would also highlight that our findings establish the fly as a powerful model for precisely exploring how the HIF1α regulates the Warburg effect. Previous studies of breast cancer cells demonstrate that ERRα is a critical component of the Hif1α transcriptional complexes that regulate expression of hypoxia-inducible genes (Ao et al., 2008). Moreover, the HIF1α-ERR interaction appears to be of notable importance in prostate cancer, where increased expression of ERRα correlates with poor survival rates (Fujimura et al., 2007), and biochemical studies suggest that elevated levels of ERRα protects Hif1α from degradation (Zou et al., 2014). The fly provides an appealing in vivo model to further study this molecular interaction in the context of the Warburg effect. Notably the precise nature by which glycolytic gene expression is activated during the embryonic metabolic transition serves as an experimental foundation to determine the molecular mechanisms that activate and stabilize Sima/HIF1α and dERR prior to activation of the Warburg effect.

Finally, our study demonstrates that the canonical *sima* allele, *sima^KG07607^*, is a hypomorphic allele. Since *sima^KG07607^*has been extensively used by the *Drosophila* community, our study raises the question as to what other functions Sima/Hif1α might also have gone unnoticed due to use of this allele. Thus, we would encourage members of the *Drosophila* community to use *sima^KG07607^* with caution in future studies.

## ACKNOWLEDGEMENTS

We thank the Bloomington *Drosophila* Stock Center (NIH P40OD018537) for providing fly stocks, the *Drosophila* Genomics Resource Center (NIH 2P40OD010949) for genomic reagents, and Flybase (NIH 5U41HG000739). These studies were also supported by the Indiana University Light Microscopy Center (LMIC) and the Indiana University Center for Genomics and Bioinformatics (CGB). L.M. was supported by a summer research grant from the Hutton Honors College at Indiana University. J.M.T. is supported by the National Institute of General Medical Sciences of the National Institutes of Health under a R35 Maximizing Investigators’ Research Award (MIRA) 1R35GM119557.

---

**Figure S1.**
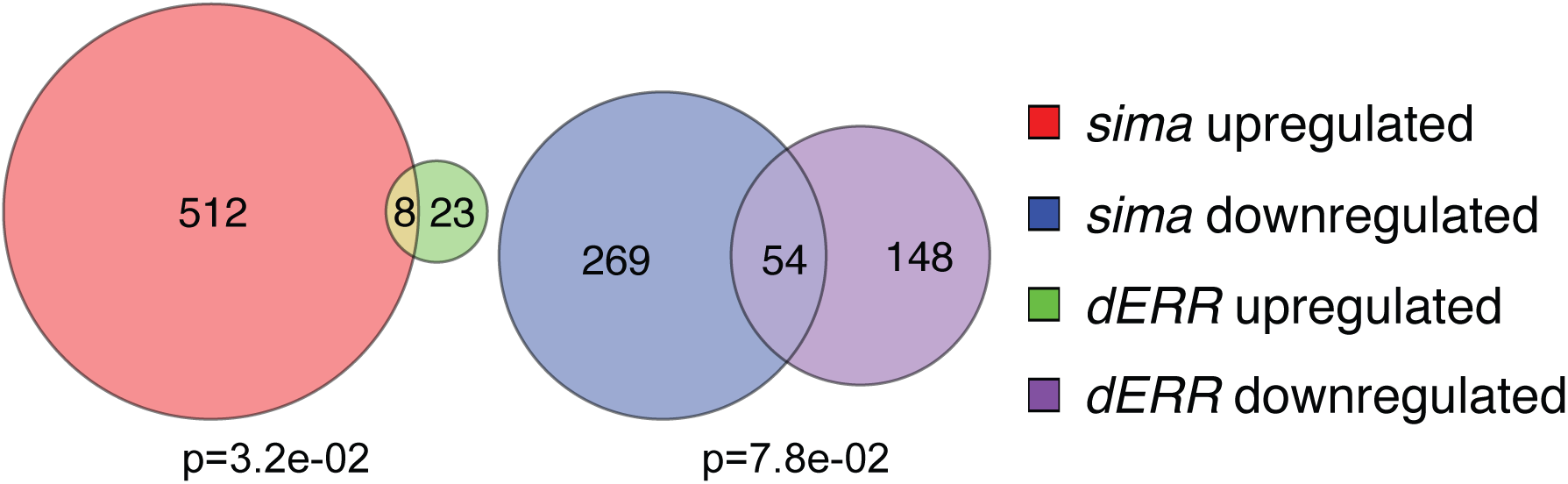
A Euler plot showing overlap from the significantly up- and down-regulated genes in RNA-seq datasets generated from *dERR* and *sima* mutant larvae (|log2 fold change|>1 and p value < .005). Significance analyzed using Fisher’s exact test.

## LITERATURE CITED

Agani, F. & Jiang, B. H. 2013. Oxygen-independent regulation of HIF-1: novel involvement of PI3K/AKT/mTOR pathway in cancer. Curr Cancer Drug Targets, 13, 245–51.

Andrews, S. 2010. FastQC: a quality control tool for high throughput sequence data. [Online]. Available: http://www.bioinformatics.babraham.ac.uk/projects/fastqc [Accessed].

Ao, A., Wang, H., Kamarajugadda, S. & Lu, J. 2008. Involvement of estrogen-related receptors in transcriptional response to hypoxia and growth of solid tumors. Proc Natl Acad Sci U S A, 105, 7821–6.

Arquier, N., Vigne, P., Duplan, E., Hsu, T., Therond, P. P., Frelin, C. & D’angelo, G. 2006. Analysis of the hypoxia-sensing pathway in Drosophila melanogaster. Biochem J, 393, 471–80.

Bacon, N. C., Wappner, P., O’rourke, J. F., Bartlett, S. M., Shilo, B., Pugh, C. W. & Ratcliffe, P. J. 1998. Regulation of the Drosophila bHLH-PAS protein Sima by hypoxia: functional evidence for homology with mammalian HIF-1 alpha. Biochem Biophys Res Commun, 249, 811–6.

Bailey, A. P., Koster, G., Guillermier, C., Hirst, E. M., Macrae, J. I., Lechene, C. P., Postle, A. D. & Gould, A. P. 2015. Antioxidant Role for Lipid Droplets in a Stem Cell Niche of Drosophila. Cell, 163, 340–53.

Baird, N. A., Turnbull, D. W. & Johnson, E. A. 2006. Induction of the heat shock pathway during hypoxia requires regulation of heat shock factor by hypoxia-inducible factor-1. J Biol Chem, 281, 38675–81.

Bandarra, D., Biddlestone, J., Mudie, S., Müller, H.-A. J. & Rocha, S. 2015. HIF-1α restricts NF-κB-dependent gene expression to control innate immunity signals. Disease Models & Mechanisms, 8, 169–181.

Bruick, R. K. & Mcknight, S. L. 2001. A conserved family of prolyl-4-hydroxylases that modify HIF. Science, 294, 1337–40.

Buddika, K., Xu, J., Ariyapala, I. S. & Sokol, N. S. 2021. I-KCKT allows dissection-free RNA profiling of adult Drosophila intestinal progenitor cells. Development, 148.

Centanin, L., Dekanty, A., Romero, N., Irisarri, M., Gorr, T. A. & Wappner, P. 2008. Cell autonomy of HIF effects in Drosophila: tracheal cells sense hypoxia and induce terminal branch sprouting. Dev Cell, 14, 547–58.

Centanin, L., Ratcliffe, P. J. & Wappner, P. 2005. Reversion of lethality and growth defects in Fatiga oxygen-sensor mutant flies by loss of hypoxia-inducible factor-alpha/Sima. EMBO Rep, 6, 1070–5.

Chatterjee, N. & Perrimon, N. 2021. What fuels the fly: Energy metabolism in Drosophila and its application to the study of obesity and diabetes. Sci Adv, 7.

Cheng, J., Kang, X., Zhang, S. & Yeh, E. T. 2007. SUMO-specific protease 1 is essential for stabilization of HIF1alpha during hypoxia. Cell, 131, 584–95.

Chowdhury, R., Yeoh, K. K., Tian, Y. M., Hillringhaus, L., Bagg, E. A., Rose, N. R., Leung, I. K., Li, X. S., Woon, E. C., Yang, M., Mcdonough, M. A., King, O. N., Clifton, I. J., Klose, R. J., Claridge, T. D., Ratcliffe, P. J., Schofield, C. J. & Kawamura, A. 2011. The oncometabolite 2-hydroxyglutarate inhibits histone lysine demethylases. EMBO Rep, 12, 463–9.

Church, R. B. & Robertson, F. W. 1966. Biochemical analysis of genetic differences in the growth of Drosophila. Genet Res, 7, 383–407.

Cockman, M. E., Masson, N., Mole, D. R., Jaakkola, P., Chang, G. W., Clifford, S. C., Maher, E. R., Pugh, C. W., Ratcliffe, P. J. & Maxwell, P. H. 2000. Hypoxia inducible factor-alpha binding and ubiquitylation by the von Hippel-Lindau tumor suppressor protein. J Biol Chem, 275, 25733–41.

Consortium, T. G. O., Aleksander, S. A., Balhoff, J., Carbon, S., Cherry, J. M., Drabkin, H. J., Ebert, D., Feuermann, M., Gaudet, P., Harris, N. L., Hill, D. P., Lee, R., Mi, H., Moxon, S., Mungall, C. J., Muruganugan, A., Mushayahama, T., Sternberg, P. W., Thomas, P. D., Van Auken, K., Ramsey, J., Siegele, D. A., Chisholm, R. L., Fey, P., Aspromonte, M. C., Nugnes, M. V., Quaglia, F., Tosatto, S., Giglio, M., Nadendla, S., Antonazzo, G., Attrill, H., DOS Santos, G., Marygold, S., Strelets, V., Tabone, C. J., Thurmond, J., Zhou, P., Ahmed, S. H., Asanitthong, P., Luna Buitrago, D., Erdol, M. N., Gage, M. C., ALI Kadhum, M., Li, K. Y. C., Long, M., Michalak, A., Pesala, A., Pritazahra, A., Saverimuttu, S. C. C., Su, R., Thurlow, K. E., Lovering, R. C., Logie, C., Oliferenko, S., Blake, J., Christie, K., Corbani, L., Dolan, M. E., Drabkin, H. J., Hill, D. P., Ni, L., Sitnikov, D., Smith, C., Cuzick, A., Seager, J., Cooper, L., Elser, J., Jaiswal, P., Gupta, P., Jaiswal, P., Naithani, S., Lera-Ramirez, M., Rutherford, K., Wood, V., DE Pons, J. L., Dwinell, M. R., Hayman, G. T., Kaldunski, M. L., Kwitek, A. E., Laulederkind, S. J. F., Tutaj, M. A., Vedi, M., Wang, S.-J., D’eustachio, P., Aimo, L., Axelsen, K., Bridge, A., HYKA-Nouspikel, N., Morgat, A., Aleksander, S. A., Cherry, J. M., Engel, S. R., Karra, K., Miyasato, S. R., Nash, R. S., Skrzypek, M. S., Weng, S., Wong, E. D., Bakker, E., et al. 2023. The Gene Ontology knowledgebase in 2023. Genetics, 224.

Dekanty, A., Lavista-Llanos, S., Irisarri, M., Oldham, S. & Wappner, P. 2005. The insulin-PI3K/TOR pathway induces a HIF-dependent transcriptional response in Drosophila by promoting nuclear localization of HIF-alpha/Sima. J Cell Sci, 118, 5431–41.

Dobin, A., Davis, C. A., Schlesinger, F., Drenkow, J., Zaleski, C., Jha, S., Batut, P., Chaisson, M. & Gingeras, T. R. 2013. STAR: ultrafast universal RNA-seq aligner. Bioinformatics, 29, 15–21.

Doronkin, S., Djagaeva, I., Nagle, M. E., Reiter, L. T. & Seagroves, T. N. 2010. Dose-dependent modulation of HIF-1alpha/sima controls the rate of cell migration and invasion in Drosophila ovary border cells. Oncogene, 29, 1123–34.

Epstein, A. C., Gleadle, J. M., Mcneill, L. A., Hewitson, K. S., O’rourke, J., Mole, D. R., Mukherji, M., Metzen, E., Wilson, M. I., Dhanda, A., Tian, Y. M., Masson, N., Hamilton, D. L., Jaakkola, P., Barstead, R., Hodgkin, J., Maxwell, P. H., Pugh, C. W., Schofield, C. J. & Ratcliffe, P. J. 2001. C. elegans EGL-9 and mammalian homologs define a family of dioxygenases that regulate HIF by prolyl hydroxylation. Cell, 107, 43–54.

Feldser, D., Agani, F., Iyer, N. V., Pak, B., Ferreira, G. & Semenza, G. L. 1999. Reciprocal positive regulation of hypoxia-inducible factor 1alpha and insulin-like growth factor 2. Cancer Res, 59, 3915–8.

Fujimura, T., Takahashi, S., Urano, T., Kumagai, J., Ogushi, T., Horie-Inoue, K., Ouchi, Y., Kitamura, T., Muramatsu, M. & Inoue, S. 2007. Increased expression of estrogen-related receptor alpha (ERRalpha) is a negative prognostic predictor in human prostate cancer. Int J Cancer, 120, 2325–30.

Gillette, C. M., Tennessen, J. M. & Reis, T. 2021. Balancing energy expenditure and storage with growth and biosynthesis during Drosophila development. Dev Biol, 475, 234–244.

Gramates, L. S., Agapite, J., Attrill, H., Calvi, B. R., Crosby, M. A., Dos Santos, G., Goodman, J. L., Goutte-Gattat, D., Jenkins, V. K., Kaufman, T., Larkin, A., Matthews, B. B., Millburn, G., Strelets, V. B. & Consortium, T. F. 2022. FlyBase: a guided tour of highlighted features. Genetics, 220.

Gratz, S. J., Ukken, F. P., Rubinstein, C. D., Thiede, G., Donohue, L. K., Cummings, A. M. & O’Connor-Giles, K. M. 2014. Highly specific and efficient CRISPR/Cas9-catalyzed homology-directed repair in Drosophila. Genetics, 196, 961–71.

Hayashi, Y., Yokota, A., Harada, H. & Huang, G. 2019. Hypoxia/pseudohypoxia-mediated activation of hypoxia-inducible factor-1alpha in cancer. Cancer Sci, 110, 1510–1517.

Hu, Y., Comjean, A., Attrill, H., Antonazzo, G., Thurmond, J., Chen, W., Li, F., Chao, T., Mohr, S. E., Brown, N. H. & Perrimon, N. 2023. PANGEA: a new gene set enrichment tool for Drosophila and common research organisms. Nucleic Acids Research, 51, W419–W426.

Intlekofer, A. M., Wang, B., Liu, H., Shah, H., Carmona-Fontaine, C., Rustenburg, A. S., Salah, S., Gunner, M. R., Chodera, J. D., Cross, J. R. & Thompson, C. B. 2017. L-2-Hydroxyglutarate production arises from noncanonical enzyme function at acidic pH. Nat Chem Biol, 13, 494–500.

Isaacs, J. S., Jung, Y. J., Mimnaugh, E. G., Martinez, A., Cuttitta, F. & Neckers, L. M. 2002. Hsp90 regulates a von Hippel Lindau-independent hypoxia-inducible factor-1 alpha-degradative pathway. J Biol Chem, 277, 29936–44.

Joshi, S., Singh, A. R. & Durden, D. L. 2014. MDM2 regulates hypoxic hypoxia-inducible factor 1alpha stability in an E3 ligase, proteasome, and PTEN-phosphatidylinositol 3-kinase-AKT-dependent manner. J Biol Chem, 289, 22785–22797.

Kaelin, W. G., JR. & Ratcliffe, P. J. 2008. Oxygen sensing by metazoans: the central role of the HIF hydroxylase pathway. Mol Cell, 30, 393–402.

Kamura, T., Sato, S., Iwai, K., Czyzyk-Krzeska, M., Conaway, R. C. & Conaway, J. W. 2000. Activation of HIF1alpha ubiquitination by a reconstituted von Hippel-Lindau (VHL) tumor suppressor complex. Proc Natl Acad Sci U S A, 97, 10430–5.

Lavista-Llanos, S., Centanin, L., Irisarri, M., Russo, D. M., Gleadle, J. M., Bocca, S. N., Muzzopappa, M., Ratcliffe, P. J. & Wappner, P. 2002. Control of the hypoxic response in Drosophila melanogaster by the basic helix-loop-helix PAS protein similar. Mol Cell Biol, 22, 6842–53.

Lee, B., Barretto, E. C. & Grewal, S. S. 2019. TORC1 modulation in adipose tissue is required for organismal adaptation to hypoxia in Drosophila. Nat Commun, 10, 1878.

Lee, J. H., Kim, E. J., Kim, D. K., Lee, J. M., Park, S. B., Lee, I. K., Harris, R. A., Lee, M. O. & Choi, H. S. 2012. Hypoxia induces PDK4 gene expression through induction of the orphan nuclear receptor ERRγ. PLoS One, 7, e46324.

Li, H., Chawla, G., Hurlburt, A. J., Sterrett, M. C., Zaslaver, O., Cox, J., Karty, J. A., Rosebrock, A. P., Caudy, A. A. & Tennessen, J. M. 2017. Drosophila larvae synthesize the putative oncometabolite L-2-hydroxyglutarate during normal developmental growth. Proc Natl Acad Sci U S A, 114, 1353–1358.

Li, H., Handsaker, B., Wysoker, A., Fennell, T., Ruan, J., Homer, N., Marth, G., Abecasis, G. & Durbin, R. 2009. The Sequence Alignment/Map format and SAMtools. Bioinformatics, 25, 2078–9.

Li, H. & Tennessen, J. M. 2017. Methods for studying the metabolic basis of Drosophila development. Wiley Interdiscip Rev Dev Biol, 6.

Li, H. & Tennessen, J. M. 2018. Preparation of Drosophila Larval Samples for Gas Chromatography-Mass Spectrometry (GC-MS)-based Metabolomics. J Vis Exp.

Li, Y., Padmanabha, D., Gentile, L. B., Dumur, C. I., Beckstead, R. B. & Baker, K. D. 2013. HIF- and non-HIF-regulated hypoxic responses require the estrogen-related receptor in Drosophila melanogaster. PLoS Genet, 9, e1003230.

Liao, Y., Smyth, G. K. & Shi, W. 2019. The R package Rsubread is easier, faster, cheaper and better for alignment and quantification of RNA sequencing reads. Nucleic Acids Res, 47, e47.

Liu, Y. V. & Semenza, G. L. 2007. RACK1 vs. HSP90: competition for HIF-1 alpha degradation vs. stabilization. Cell Cycle, 6, 656–9.

Loenarz, C., Coleman, M. L., Boleininger, A., Schierwater, B., Holland, P. W., Ratcliffe, P. J. & Schofield, C. J. 2011. The hypoxia-inducible transcription factor pathway regulates oxygen sensing in the simplest animal, Trichoplax adhaerens. EMBO Rep, 12, 63–70.

Love, M. I., Huber, W. & Anders, S. 2014. Moderated estimation of fold change and dispersion for RNA-seq data with DESeq2. Genome Biol, 15, 550.

Magar, A. G., Morya, V. K., Kwak, M. K., Oh, J. U. & Noh, K. C. 2024. A Molecular Perspective on HIF-1alpha and Angiogenic Stimulator Networks and Their Role in Solid Tumors: An Update. Int J Mol Sci, 25.

Martin, M. 2011. Cutadapt removes adapter sequences from high-throughput sequencing reads. 2011, 17, 3.

Maxwell, P. H., Wiesener, M. S., Chang, G. W., Clifford, S. C., Vaux, E. C., Cockman, M. E., Wykoff, C. C., Pugh, C. W., Maher, E. R. & Ratcliffe, P. J. 1999. The tumour suppressor protein VHL targets hypoxia-inducible factors for oxygen-dependent proteolysis. Nature, 399, 271–5.

Mortimer, N. T. & Moberg, K. H. 2009. Regulation of Drosophila embryonic tracheogenesis by dVHL and hypoxia. Dev Biol, 329, 294–305.

Mortimer, N. T. & Moberg, K. H. 2013. The archipelago ubiquitin ligase subunit acts in target tissue to restrict tracheal terminal cell branching and hypoxic-induced gene expression. PLoS Genet, 9, e1003314.

Mukherjee, T., Kim, W. S., Mandal, L. & Banerjee, U. 2011. Interaction between Notch and Hif-alpha in development and survival of Drosophila blood cells. Science, 332, 1210–3.

Nadtochiy, S. M., Schafer, X., Fu, D., Nehrke, K., Munger, J. & Brookes, P. S. 2016. Acidic pH Is a Metabolic Switch for 2-Hydroxyglutarate Generation and Signaling. J Biol Chem, 291, 20188–97.

Nambu, J. R., Chen, W., Hu, S. & Crews, S. T. 1996. The Drosophila melanogaster similar bHLH-PAS gene encodes a protein related to human hypoxia-inducible factor 1 alpha and Drosophila single-minded. Gene, 172, 249–54.

Ostberg, T., Jacobsson, M., Attersand, A., Mata de Urquiza, A. & Jendeberg, L. 2003. A triple mutant of the Drosophila ERR confers ligand-induced suppression of activity. Biochemistry, 42, 6427–35.

Öztürk-Çolak, A., Marygold, S. J., Antonazzo, G., Attrill, H., Goutte-Gattat, D., Jenkins, V. K., Matthews, B. B., Millburn, G., Dos Santos, G., Tabone, C. J. & Consortium, T. F. 2024. FlyBase: updates to the Drosophila genes and genomes database. Genetics.

Pang, Z., Lu, Y., Zhou, G., Hui, F., Xu, L., Viau, C., Spigelman, Aliya F., Macdonald, Patrick E., Wishart, David S., Li, S. & Xia, J. 2024. MetaboAnalyst 6.0: towards a unified platform for metabolomics data processing, analysis and interpretation. Nucleic Acids Research, 52, W398–W406.

Ravi, R., Mookerjee, B., Bhujwalla, Z. M., Sutter, C. H., Artemov, D., Zeng, Q., Dillehay, L. E., Madan, A., Semenza, G. L. & Bedi, A. 2000. Regulation of tumor angiogenesis by p53-induced degradation of hypoxia-inducible factor 1alpha. Genes Dev, 14, 34–44.

Romero, N. M., Dekanty, A. & Wappner, P. 2007. Cellular and developmental adaptations to hypoxia: a Drosophila perspective. Methods Enzymol, 435, 123–44.

Romero, N. M., Irisarri, M., Roth, P., Cauerhff, A., Samakovlis, C. & Wappner, P. 2008. Regulation of the Drosophila hypoxia-inducible factor alpha Sima by CRM1-dependent nuclear export. Mol Cell Biol, 28, 3410–23.

Sonnenfeld, M., Ward, M., Nystrom, G., Mosher, J., Stahl, S. & Crews, S. 1997. The Drosophila tango gene encodes a bHLH-PAS protein that is orthologous to mammalian Arnt and controls CNS midline and tracheal development. Development, 124, 4571–82.

Tamamouna, V., Rahman, M. M., Petersson, M., Charalambous, I., Kux, K., Mainor, H., Bolender, V., Isbilir, B., Edgar, B. A. & Pitsouli, C. 2021. Remodelling of oxygen-transporting tracheoles drives intestinal regeneration and tumorigenesis in Drosophila. Nat Cell Biol, 23, 497–510.

Tanimoto, K., Makino, Y., Pereira, T. & Poellinger, L. 2000. Mechanism of regulation of the hypoxia-inducible factor-1 alpha by the von Hippel-Lindau tumor suppressor protein. EMBO J, 19, 4298–309.

Tennessen, J. M., Baker, K. D., Lam, G., Evans, J. & Thummel, C. S. 2011. The Drosophila estrogen-related receptor directs a metabolic switch that supports developmental growth. Cell Metab, 13, 139–48.

Tennessen, J. M., Barry, W. E., Cox, J. & Thummel, C. S. 2014a. Methods for studying metabolism in Drosophila. Methods, 68, 105–15.

Tennessen, J. M., Bertagnolli, N. M., Evans, J., Sieber, M. H., Cox, J. & Thummel, C. S. 2014b. Coordinated metabolic transitions during Drosophila embryogenesis and the onset of aerobic glycolysis. G3 (Bethesda), 4, 839–50.

Texada, M. J., Jørgensen, A. F., Christensen, C. F., Koyama, T., Malita, A., Smith, D. K., Marple, D. F. M., Danielsen, E. T., Petersen, S. K., Hansen, J. L., Halberg, K. A. & Rewitz, K. F. 2019. A fat-tissue sensor couples growth to oxygen availability by remotely controlling insulin secretion. Nature Communications, 10, 1955.

Vasilia, T. & Chrysoula, P. 2018. The Hypoxia-Inducible Factor-1α in Angiogenesis and Cancer: Insights from the Drosophila Model. In: Fumiaki, U. (ed.) Gene Expression and Regulation in Mammalian Cells. Rijeka: IntechOpen.

Venken, K. J., Carlson, J. W., Schulze, K. L., Pan, H., He, Y., Spokony, R., Wan, K. H., Koriabine, M., De Jong, P. J., White, K. P., Bellen, H. J. & Hoskins, R. A. 2009. Versatile P[acman] BAC libraries for transgenesis studies in Drosophila melanogaster. Nat Methods, 6, 431–4.

Vigne, P. & Frelin, C. 2008. The role of polyamines in protein-dependent hypoxic tolerance of Drosophila. BMC Physiology, 8, 22.

Wang, C. W., Purkayastha, A., Jones, K. T., Thaker, S. K. & Banerjee, U. 2016. In vivo genetic dissection of tumor growth and the Warburg effect. Elife, 5.

Zou, C., Yu, S., Xu, Z., Wu, D., Ng, C. F., Yao, X., Yew, D. T., Vanacker, J. M. & Chan, F. L. 2014. ERRα augments HIF-1 signalling by directly interacting with HIF-1α in normoxic and hypoxic prostate cancer cells. J Pathol, 233, 61–73.

